# HyPo: Super Fast & Accurate Polisher for Long Read Genome Assemblies

**DOI:** 10.1101/2019.12.19.882506

**Authors:** Ritu Kundu, Joshua Casey, Wing-Kin Sung

**Affiliations:** Department of Computer Science, National University of Singapore, Singapore 117417; Department of Computational and Systems Biology, Genome Institute of Singapore, Singapore 138672

**Keywords:** Polishers, Long read assemblies, Human genomes, Hypo

## Abstract

Efforts towards making population-scale long read genome assemblies (especially human genomes) viable have intensified recently with the emergence of many fast assemblers. The reliance of these fast assemblers on polishing for the accuracy of assemblies makes it crucial. We present HyPo–a **Hy**brid **Po**lisher–that utilises short as well as long reads within a single run to polish a long read assembly of small and large genomes. It exploits unique genomic kmers to selectively polish segments of contigs using partial order alignment of selective read-segments. As demonstrated on human genome assemblies, Hypo generates significantly more accurate polished assemblies in about one-third time with about half the memory requirements in comparison to Racon (the widely used polisher currently).

## 1 Introduction

Genome assembly–reconstructing a genome from the fragments (reads) produced by a DNA sequencer–and its analysis to explore genetic variations amongst or within species is central to genomics. Third-generation (DNA) sequencers (TGS) like Pacific Biosciences (PacBio) and Oxford Nanopore Technologies (ONT) have given new impetus to genomics by enabling high quality assembly and analysis of genomes. Overcoming the major limitation of second-generation or next-generation sequencers (NGS; e.g. Illumina) stemming from read-length being limited to several hundred base pairs, TGS produce reads with an average length of tens of thousands of base pairs which leads to more contiguous assembly, resolution of more repetitive elements, and revelation of larger structural variations [Roberts et al., 2013, Lee et al., 2016]. Till very recently, *de novo* assembly of human genome was done using the ultra-long-reads from nanopore (along with other complimentary technologies to improve the quality) which not only exceeds the continuity of the human reference genome (GRCH38) but also produces a telomere-to-telomere complete Chromosome X [Miga et al., 2019]. However, in contrast to the NGS short reads with accuracy greater than 99%, long reads suffer from the drawback of higher error rates (> 10%) [Weirather et al., 2017, Jain et al., 2018]. Moreover, the error profile of long reads skews towards insertion-deletions (indels) than substitutions; homopolymer indels being more prominent amongst those [Weirather et al., 2017].

To deal with noisy long reads, assemblers typically resort to correcting errors before assembly which is computationally expensive, especially for large genomes [Sović et al., 2016, Fu et al., 2019, Zhang et al., 2019]. In addition, assembled contigs are polished–base accuracy improvement by using consensus generation from the reads–in the end to enhance the assembly quality. Recently, several fast assemblers (miniasm [Li, 2016], Ra [Vaser and Šikić, 2019], wtdbg2 [Ruan and Li, 2019]) have been made available which circumvent the resource and time intensive step of read error correction and thereby assemble structurally correct contigs an order of magnitude faster, albeit with more base level errors (about 10 times more errors as compared to those with other assemblers which make use of corrected reads). These fast assemblers rely solely on polishing for error-correction. Importantly, in long read assemblies, it is critical to correct errors in order to avoid their grave impact on protein prediction owing to the higher error rate and indel predominance [Watson and Warr, 2019]. Thus, polishing tools play a crucial role in producing accurate long read assemblies, especially the ones generated by fast, error-correction-free assemblers.

Polishers can broadly be categorised as ‘Sequencer-bound’ and ‘General’. Sequencer-bound polishers require raw signal-level information generated by a particular sequencer and therefore those can polish using reads only from a specific sequencer. For ONT, NanoPolish [Loman et al., 2015] and its successor Medaka [Nanopore Technologies, 2019] fall into this category. Similarly, for PacBio, Quiver [Chin et al., 2013] and its successor Arrow [Laird Smith et al., 2016] are available.

General polishers, on the other hand, are robust enough to work with reads generated by any sequencer. Earlier, Pilon [Walker et al., 2014] has been a widely used general polisher for bacterial and small (< 100 bp long) eukaryotic genomes. However, Pilon is now increasingly being supplanted by (or used in combination with) Racon [Vaser et al., 2017] owing to the ultra-fast speed of the latter due to which it scales well resource-wise on large genomes. Several new polishers have emerged more recently: wtpoa-cns (stand alone consensus module of wtdbg2), ntEdit [Warren et al., 2019], Apollo [Firtina et al., 2019]. Each polisher other than ntEdit rely on the alignment information of reads on the draft (uncorrected) assembly. Pilon is based on the pileup of bases from the reads at each base position in the draft contigs. Racon and wtpoa-cns use consensus generation from graphical Partial Order Alignment (POA) [Lee et al., 2002, Lee, 2003] of multiple sequences (i.e. of aligned read-segments). Both Racon and wtpoa-cns (conceptually) break contigs into smaller segments and use a single instruction multiple data (SIMD) implementation of POA in order to make it faster and thus practical. Apollo deploys a machine-learning approach to build a profile Hidden Markov Model (pHMM) [Firtina et al., 2018] of the draft assembly which is then used to correct errors. In contrast to exploiting read-to-assembly alignments, ntEdit corrects error based on scanning kmers (sequences of length k) in the draft and checking their presence/absence utilising Bloom Filters that store kmers in the reads.

Each general polisher has its own share of limitations. Pilon and ntEdit are designed to work primarily with short reads which are remarkably accurate. Additionally, as mentioned above, Pilon does not scale well on large genomes in terms of resources.On the other hand, Racon and wtpoa-cns are fast and scalable on large genomes and can handle short as well as noisy long reads. However, in one run, either only long or only short reads can be used for polishing by both; long read polishing followed by short read polishing is recommended for better accuracy. Apollo can use both types of reads within a single run but it is excruciatingly slow; for instance, it took about two and a half hours to polish an E.Coli data-set using PacBio reads where Racon took only about 2 minutes [Firtina et al., 2019]. Currently, Racon is the widely used polisher given its speed and relatively better accuracy (our results also confirm that Racon, overall, produces more accurate results than other polishers).

Here, we present HyPo–a **Hy**brid **Po**lisher–that utilises short as well as long reads within a single run to polish a long reads assembly of small and large genomes. It exploits unique genomic kmers to selectively polish segments of contigs using POA of selective read-segments. We demonstrate that Hypo generates significantly more accurate polished assembly in about one-third of the time with only about half the memory requirements in comparison to Racon.

While the performance of polishers can be compared by comparing the resources (memory and time) used, the evaluation of the accuracy of polishers usually follows the same approach as the one used to compare assemblers which typically involves comparisons based on one or more of the following: similarity of contigs with the corresponding reference genome, assembly statistics (N50, % reference aligned etc.), rates of substitution/insertions/deletions. However, these accuracy assessment criteria suffer from two drawbacks: (i) Polished assemblies are not that drastically different from each other as draft assemblies which makes it difficult for these parameters to highlight differences amongst them. (ii) These parameters do not sufficiently capture the differences between the improvement of base-level accuracy by various polishers. We also present here a novel way to compare base-level accuracy of polishers if given the information about the true variants in the assembled genome. Additionally, we compare polishers on more reliable criteria to compare polished assemblies such as: (i) BUSCO Score [Simão et al., 2015] using the eukaryote set of orthologs (ii) the number of known genomic features found like genes, CDS (coding sequence), exons etc.

## 2 Methods

Broadly, we (conceptually) divide a draft (uncorrected) contig into two types of regions (segments): *Strong* and *Weak*. Strong regions are those which have strong evidence (*support*) of their correctness and thus do not need polishing. Weak regions, on the other hand, will be polished using POA. Each weak region will be polished using either short reads or long reads; short reads taking precedence over long reads. To identify strong regions, we make use of *solid* kmers (expected unique genomic kmers). Strong regions also play a role in selecting the read-segments to polish their neighbouring weak regions. Furthermore, our approach takes into account that the long reads and thus the assemblies generated from them are prone to homopolymer errors as mentioned in the beginning.

### 2.1 Identifying Strong Regions

#### Identifying solid kmers

Frequency distribution is obtained from kmer counting in the short (accurate) reads. A frequency range is determined such that the kmers with low frequency (likely errors) and those with high frequency (likely originating in repeat regions) are excluded. The kmers appearing in the selected frequency range (which is spread around read-coverage) are likely to be the kmers appearing uniquely in the genome that has been assembled. From the kmers thus extracted, we filter out those which have *homopolymer runs* (a sequence with repetition of the same base) at the either ends; we refer to the remaining kmers as **solid kmers**.

#### Finding support

We scan each contig and record the solid kmers appearing in them; considering only those occurrences that do not begin or end in homopolymers i.e. the base previous (next) to the beginning (end) of the occurrence should be different from the first (last) base of the solid kmer. Since the draft has significant base-level errors, many of these solid kmers found in contigs will be erroneous. To identify a non-erroneous solid kmer in a contig, we employ the evidence present in the reads aligned to the corresponding segment of the contig containing the kmer in the following way: If a solid kmer appears in a ‘reasonable’ distance of expected position in an aligned read, we say that the read *supports* that solid kmer; if ‘sufficient’ number of aligned reads support a solid kmer, we call it a **supported kmer**.

#### Strong regions

We (conceptually) merge the overlapping and adjacent supported kmers into a strong region avoiding those whose inclusion may have errors. The regions interspersed between strong regions become the weak regions.

### 2.2 Polishing Weak Regions

We divide the weak regions into smaller **windows** making use of minimizers [Roberts et al., 2004] which have sufficient *support* (defined in the same way as in solid kmers). Next, we try to find read-segments aligned to these windows for getting consensus using their POA. We will refer to the read-segments selected to polish a window as **arms**. Note that we may have three kinds of arms: (1) An *Internal* arm spans the whole window (read-alignment starts before and ends after the window); (2) A *Prefix* arm starts at the beginning of window but terminates without reaching the end of the window (read-alignment starts before and ends in the window); (3) A *suffix* arm starts in the window and terminates with its end (read-alignment starts in the window and ends after it). Due to the small size of a window, we will not have many arms which are completely contained within a window; if there are any, we ignore it. Before we proceed, we would like to clarify that this step bears similarity to Racon in the sense that we also divide regions to be polished into smaller windows and use alignment information to find arms for POA. However, the window-division method as well as which and how arms are being selected are substantially different.

#### Finding short arms

We use the first and the last supported kmers of the preceding (if any) and the following (if any) strong regions or preceding and following minimizers (if any) adjacent to a window as **anchors** to select the read-segments to polish the window as follows: Each non-ambiguously and primarily aligned read that has the anchors within a reasonable distance from the expected position is selected and the sequence between markers is used as the arm if it is sufficiently long. If a window has no adjacent strong region (or minimizers) at either end then we rely only on alignment information from CIGAR string to extract coordinates of the read corresponding to the window’s starting/ending coordinates. If a window has sufficient number of internal arms then its prefix/suffix arms are not used for POA consensus.

#### Finding long arms

Next, we find arms from long reads but only in those windows which do not have sufficient number of short arms. We do not seek help from neighbouring strong regions here and utilise only CIGAR string to extract segments from only those reads which are ‘sufficiently similar’ to the draft sequence in the window.

#### POA consensus

We use modified SPOA library (stand alone POA module of Racon) to derive consensus of arms in each window and replace the window sequence (draft) with the consensus sequence. We use the global alignment method for internal arms but a (customised) prefix/suffix alignment for prefix/suffix arms ensuring that the alignment starts/ends at the beginning/ending of the window. We perform two rounds of POA consensus generation in the windows with long arms.

The final polished sequence is then obtained by concatenating the polished sequence of each window/strong region/minimizer (where polished sequence of a strong region/minimizer is same as the draft sequence.)

## 3 Evaluation

We compared the results of polishing with only short reads and two-rounds polishing (first with long reads followed by that with short reads). We evaluated^i^ Racon [Vaser et al., 2017], wtpoa-cns [Ruan and Li, 2019], ntEdit [Warren et al., 2019], and Pilon [Walker et al., 2014] against Hypo for short reads only polishing. ntEdit and Pilon were excluded from the 2-rounds polishing as they do not operate with long reads. The alignment files required by most of the polishers were generated using Minimap2 [Li, 2018].

### 3.1 Data-sets

The data-sets for which evaluation results have been included in this manuscript are the whole human genome data-set HG002 (from Genome-in-a-Bottle (GIAB) [Zook et al., 2016]) and data-sets extracted from it for Chromosome 21 (Chr21) and 1 (Chr1). The preliminary results for other data-sets (not included here) show similar trends. The data-set consists of PacBio reads (69X coverage) and paired-end Illumina reads (2*250; 55X coverage). Chr21 and Chr1 have been assembled using Wtdbg2 and Canu [Koren et al., 2017] (a widely used assembler which uses read-errors correction before assembly) while the whole genome was assembled using Wtdbg2 only.

### 3.2 Accuracy Evaluation

We used GRCh38 as the reference for evaluation. For accuracy evaluation we used Quast [Mikheenko et al., 2018] to compute the BUSCO score and find the genomic features (like genes, CDS (coding sequence), exons). When we had the information about the true variants–gold standard High Confidence (HC) small variants–for the genome under consideration, we assessed the base-level errors in an assembly. We used a novel method of evaluating accuracy where computing errors takes into account the hetrozygosity of true variants. We mapped the contigs to the reference using Minimap2 and made use of paftools.js script (included in the Minimap2 package) setting up the parameters such that every difference between contigs and the reference is reported as a variant. We considered only the regions where such variants have been called for each assembly being compared and which overlapped with the region for which true variants are available. Any variant which was not listed amongst the true variants was considered an *error*. Every true variant with each haplotype being different from the reference and not listed in the called variants from the assembly was considered a *missed* variant. We compared the error variants and missed variants of each assembly.

### 3.3 Performance Evaluation

Each polisher (except Racon) was run with 48 threads on a machine running Ubuntu (18.04.2 LTS) with the following specifications: 512 GB RAM (DDR4), 48 cores (Intel(R) Xeon(R) Silver 4116 CPU @ 2.10GHz), 3.84TB SSD as the scratch. For the whole genomes, Racon could not work on the 512G machine even after trying the wrapper to split target and using the reads in fasta format rather than fastq. The performance measures for the whole genome polishing by Racon is on another machine with 8TB RAM.

## 4 Results

The following keys have been used for short reads only polishing: **r** (Racon polished), **w** (wtpoa-cns polished), **h** (Hypo polished), **n** (ntEdit polished), **p** (Pilon polished). The 2-rounds polished results by Racon and wtpoa-cns using long reads and short reads have been labelled as **r2** and **w2**, respectively. **h2** has been used to label the polished results of Hypo using both long and short reads.

### 4.1 Performance comparison

Tables 1 and 2, respectively, record the timings and the memory consumption of various polishers.

**Table 1:** Comparison of time ([hh:]mm:ss) consumed by various polishers. *W* indicates Wtdbg2 assembly and *C* indicates Canu assembly. For 2-rounds polishing, Plus sign (+) has been used as the delimiter. Colour code: Dark green, light green, and red colours have been used to highlight the best, the next best and the worst polisher w.r.t time-requirements.

|  | r | w | h | n | p | r2 | w2 | h2 |
| --- | --- | --- | --- | --- | --- | --- | --- | --- |
| Chr21 (W) | 6:52 | 4:09.35 | 2:07 | 1:44 | 7:29 | 01:22 + 6:39 | 08:04 + 4:09 | 05:29 |
| Chr21 (C) | 7:07 | 4:15.55 | 2:12 | 1:49 | 4:25 | 01:30 + 6:43 | 03:14 + 4:15 | 02:55 |
| Chr1 (W) | 47:35 | 26:35 | 12:05 | 9:24 | 27:37 | 07:38 47:56 | 51:36 26:43 | 19:17 |
| Chr1 (C) | 46:33 | 27:17 | 10:53 | 9:43 | 19:03 | 08:19 + 49:20 | 19:53 27:10 | 12:58 |
| Whole genome (W) | 12:44:33 | 06:37:49 | 2:59:18 | 2:18:32 | - | 1:21:57 + 13:13:55 | 10:48:17 + 6:27:49 | 4:12:44 |

**Table 2:** Comparison of memory (GB) consumed by various polishers. *W* indicates Wtdbg2 assembly and *C* indicates Canu assembly. For 2-rounds polishing, Comma (,) has been used as the delimiter. Colour code: Dark green, light green, and red colours have been used to highlight the best, the next best and the worst polisher w.r.t time-requirements.

|  | r | w | h | n | p | r2 | w2 | h2 |
| --- | --- | --- | --- | --- | --- | --- | --- | --- |
| Chr21 (W) | 13.95 | 0.96 | 11.66 | 0.53 | 91.9 | 8.26, 13.95 | 6.38, 0.95 | 12.74 |
| Chr21 (C) | 13.94 | 1 | 11.66 | 0.53 | 87.86 | 3.09, 13.94 | 5.12, 1.01 | 11.66 |
| Chr1 (W) | 86.84 | 1.07 | 48.07 | 2.42 | 137.26 | 44.62, 86.84 | 10.15, 1.07 | 61.79 |
| Chr1 (C) | 87.02 | 1.08 | 33.31 | 2.42 | 131.38 | 15.31, 87 | 6.27, 1.08 | 34.68 |
| Whole genome (W) | 1147.9 | 2.18 | 376.3 | 42.57 | - | 548.95 , 1147.91 | 10.78 , 1.87 | 410.08 |

For polishing using only short reads, ntEdit was the fastest polisher amongst all. However, Hypo was not much slower for all practical purposes. Racon seemed to be the slowest on small data-sets. However, Pilon (which polishes with only Illumina reads) could not be run on the whole data-set (owing to its memory requirements). In particular, Hypo took less than one-third the time as that of Racon and less than half the time of wtpoa-cns.

For 2-rounds polishing, Wtdbg2 and Racon both seemed to take similar time. Hypo completed in less than one-third the time as that of the other 2 polishers. Racon was quite fast when polishing with only long reads but extremely slow with short reads.

As for the memory requirements, Wtpoa-cns was consistent in using almost marginal RAM. ntEdit expectedly did not need much memory to run as it uses bloom filters to store kmers in the reads and only scans the contigs. Pilon was the most expensive in this context; so much so that it became infeasible to run Pilon on the whole genome. Racon also had very steep memory requirements; even the efforts of batch processing and converting reads from fastq to fasta format to reduce the file size were not enough to run Racon on the whole genome on a 512GB RAM machine. Hypo, in comparison to Racon, had only half the memory demands that could be met by a usual server machine. In addition, Hypo provides for adjusting the number of contigs that it processes in one batch which can be used to run it on smaller machines, albeit with compromising on speed.

### 4.2 Comparison of Errors

Figures 4.3 through 4.3 demonstrate the errors and the missed variants int the contigs polished by various polishers. Errors wise, Hypo undoubtedly beat the other polishers by a great margin. The second best polisher was Racon which, in comparison to Hypo, had more than three fold increase in InDel errors and about 50% increase in SNP errors. Similarly, Hypo missed the lowest number of variants amongst all polishers. Racon, the next best polisher, missed almost double the variants as that of Hypo. Adding long reads, further improved the errors in Hypo results. Interestingly, the errors in Racon polished results remained the same. Canu assembly, as expected, contained fewer errors as compared to WTDBG2 assembly (because Canu employs error correction of reads before assembling them). Consequently, the same pattern followed in the polished results.

**Figure 1:**
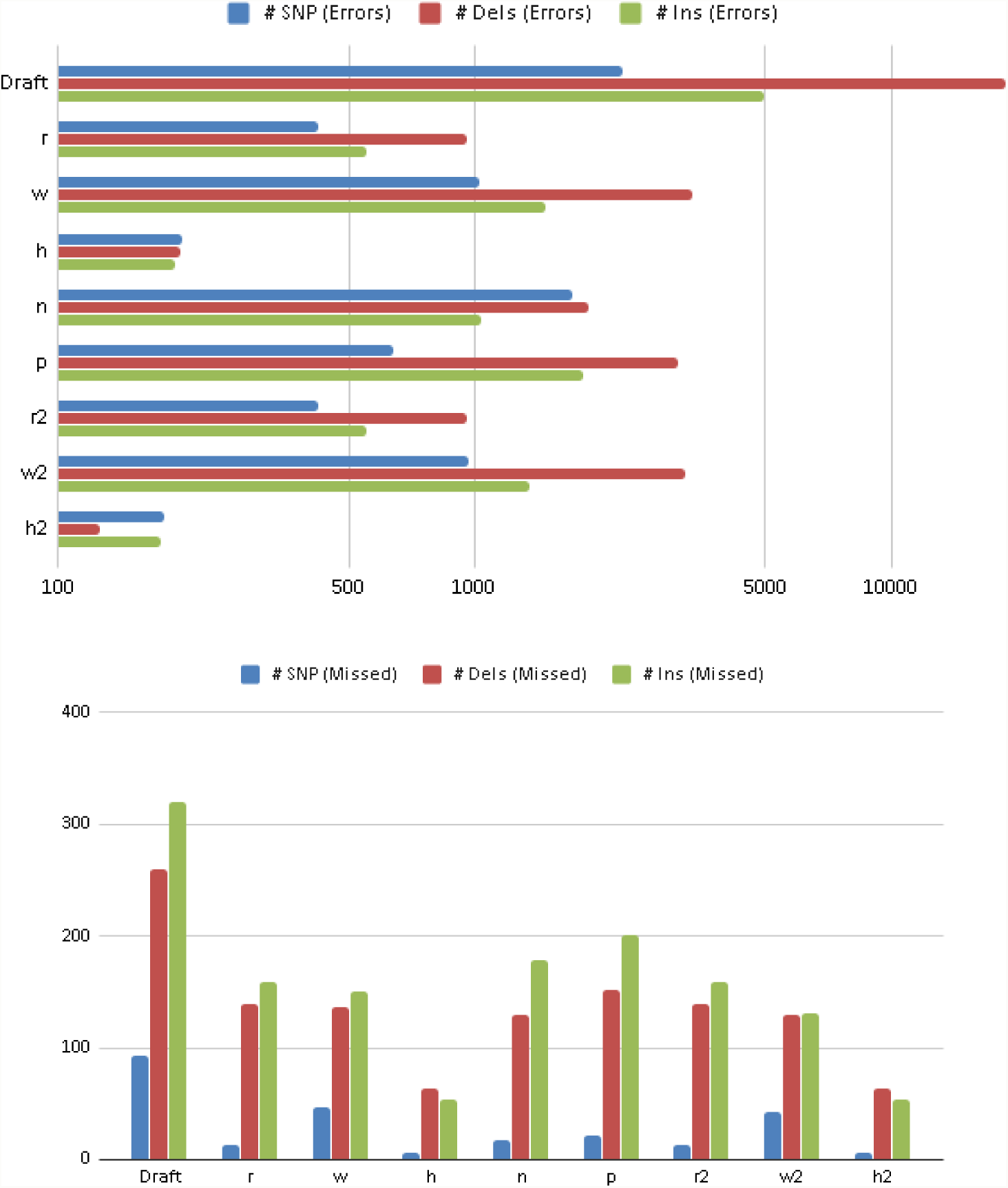
Number of errornuous (log-scale) and missed variants: Chr21 (Wtdbg2 assembly)

**Figure 2:**
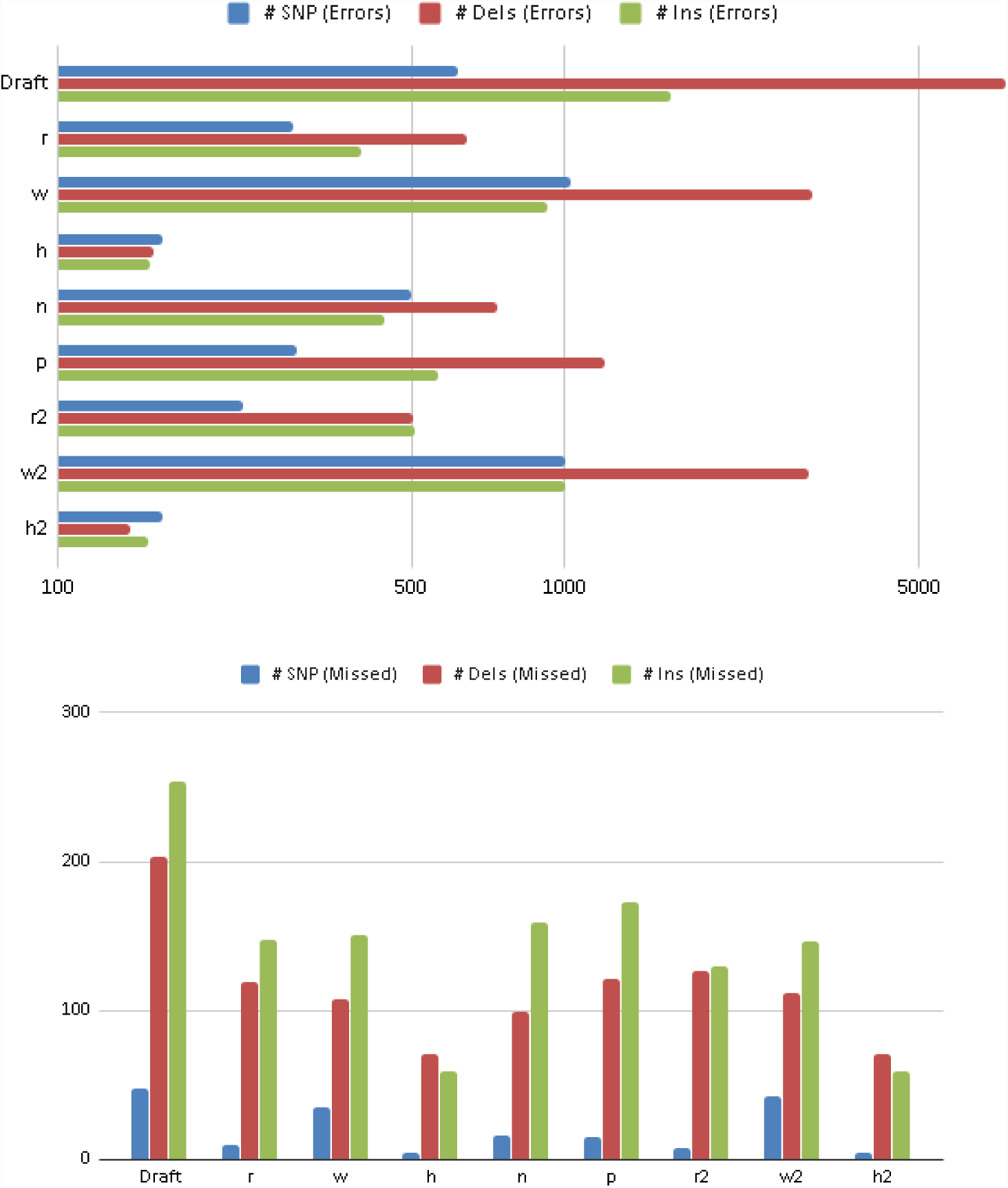
Number of errornuous (log-scale) and missed variants: Chr21 (Canu assembly)

**Figure 3:**
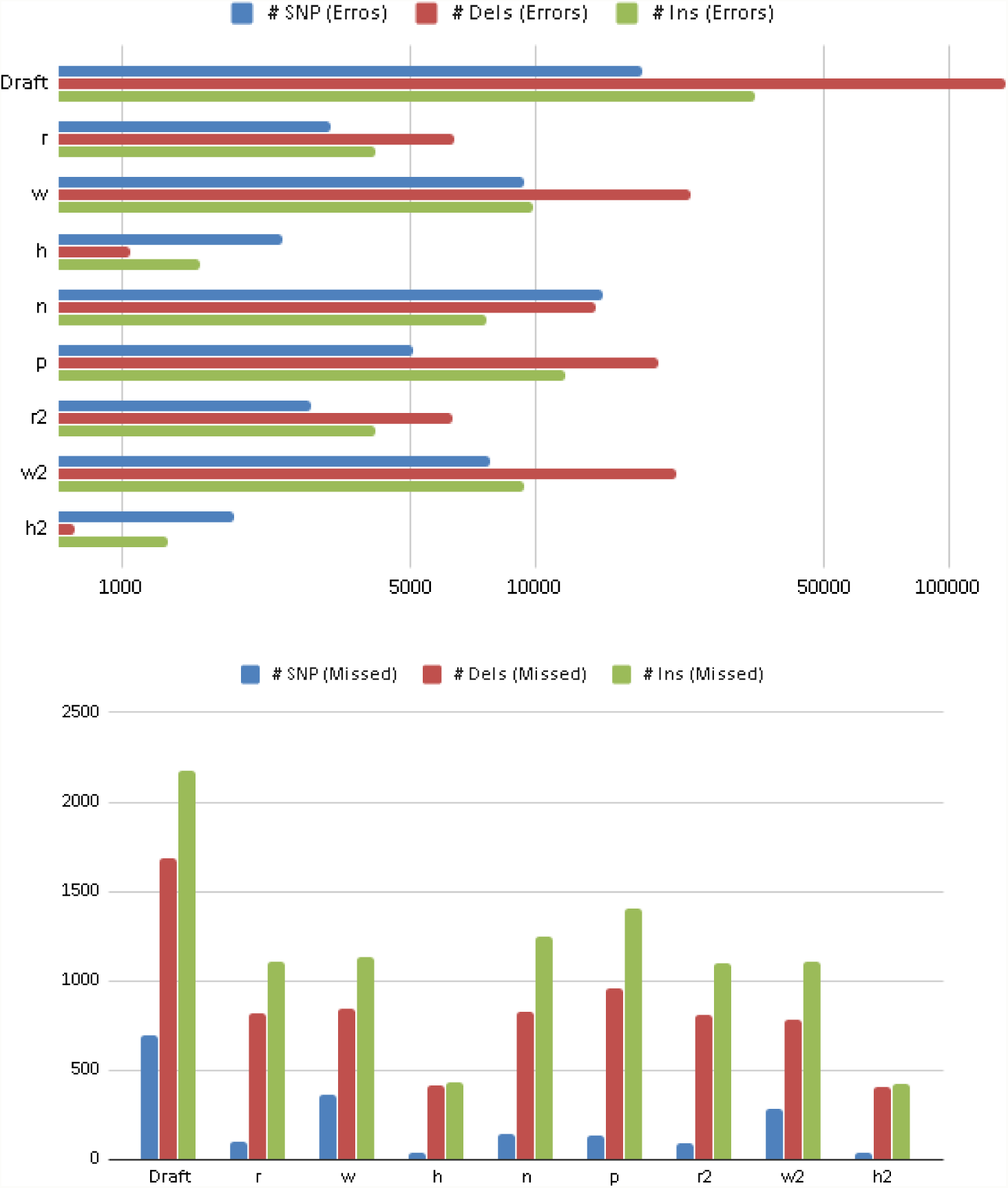
Number of errornuous (log-scale) and missed variants: Chr1 (Wtdbg2 assembly)

**Figure 4:**
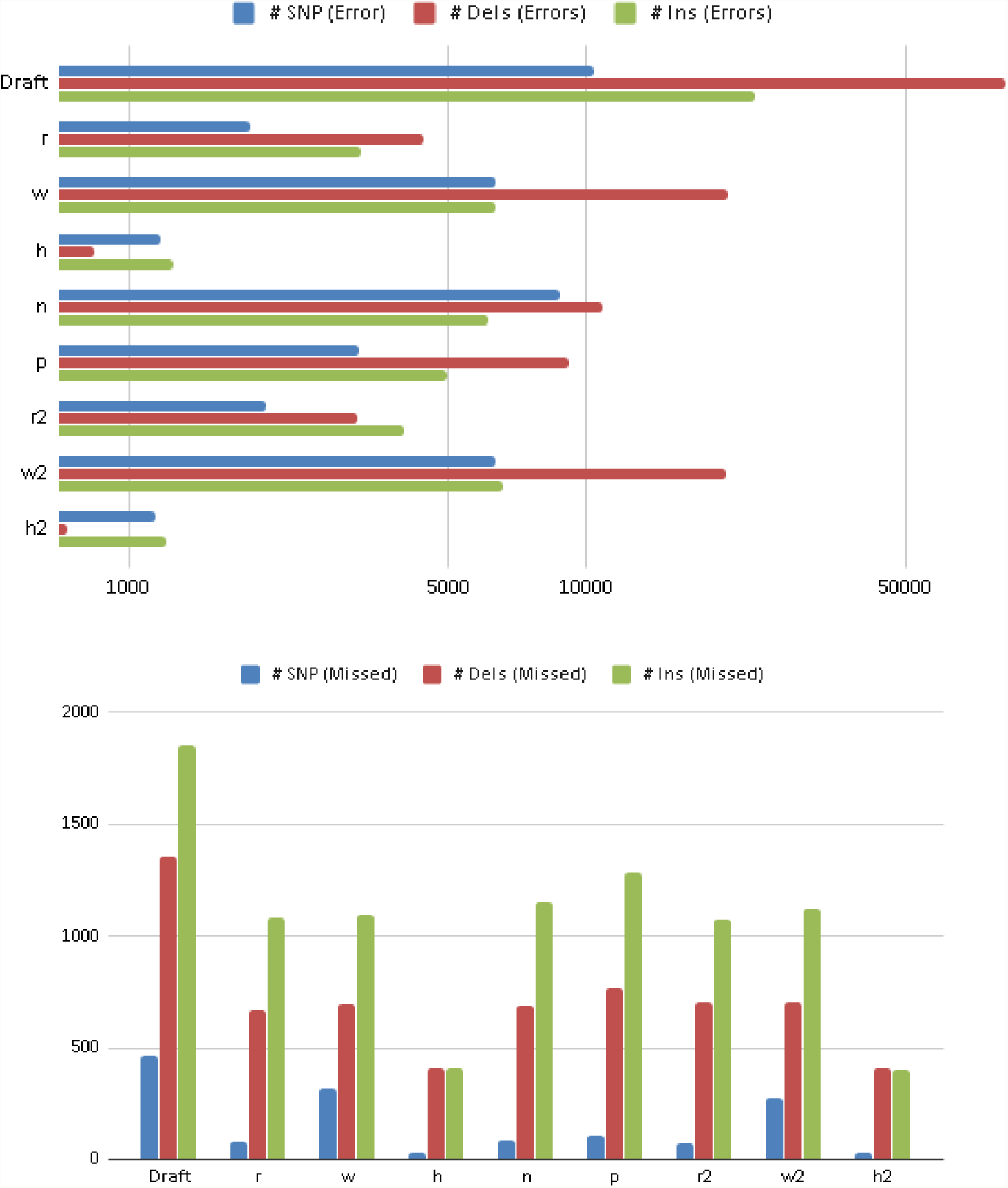
Number of errornuous (log-scale) and missed variants: Chr1 (Canu assembly)

**Figure 5:**
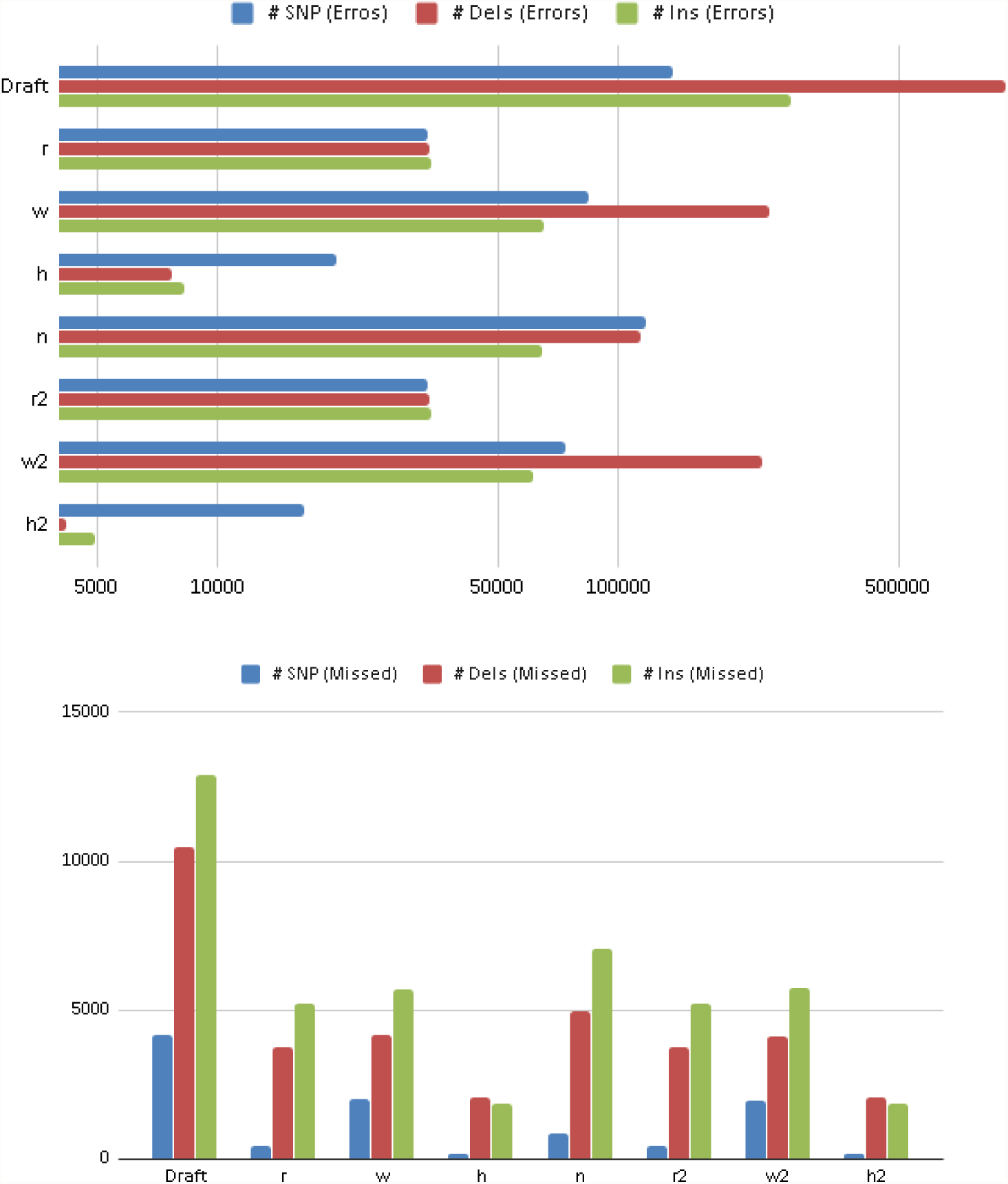
Number of errornuous (log-scale) and missed variants: Whole genome (Wtdbg2 assembly)

### 4.3 Comparison of Genomic Features and BUSCO Scores

Comparison on the basis of the number of (full) genomic features found int he contigs and the BUSCO score have been shown in Tables 3 and 4, respectively.

**Table 3:** Comparison of the genomic features found in the draft and various polished assemblies. *W* indicates Wtdbg2 assembly and *C* indicates Canu assembly. The draft features are in absolute numbers whereas the numbers for different polishers indicate the change w.r.t draft (- signs indicates the decrease). Colour code: Dark green, light green, and red colours have been used to highlight the best, the next best and the worst polisher w.r.t time-requirements.

|  | draft | r | w | h | n | p | r2 | w2 | h2 |
| --- | --- | --- | --- | --- | --- | --- | --- | --- | --- |
| Chr21 (W) | 36,478 | 655 | -257 | 689 | 579 | 618 | 655 | -422 | 732 |
| Chr21 (C) | 37,248 | 93 | -1,064 | 180 | 105 | 128 | 49 | -980 | 164 |
| Chr1 (W) | 340,581 | 2,622 | -4,488 | 2,825 | 1,682 | 2,643 | 2,622 | -3,003 | 4,195 |
| Chr1 (C) | 351,426 | 287 | -6,468 | 547 | 454 | 554 | 330 | -6,599 | 566 |
| Whole genome (W) | 3,242,831 | 85,177 | -42,916 | 96,260 | 74,984 | - | 85,177 | -32,364 | 104,096 |

**Table 4:** Comparison of the genomic features found in the draft and various polished assemblies. *W* indicates Wtdbg2 assembly and *C* indicates Canu assembly. Semicolon (;) has been used as the delimiter. Colour code: Dark green, light green, and red colours have been used to highlight the best, the next best and the worst polisher w.r.t time-requirements.

|  | draft | r | w | h | n | p | r2 | w2 | h2 |
| --- | --- | --- | --- | --- | --- | --- | --- | --- | --- |
| Chr1 (W) | 11.55;<br>1.65 | 11.88;<br>2.31 | 10.89;<br>2.31 | 12.21;<br>1.65 | 11.55;<br>1.98 | 12.21;<br>1.98 | 11.88;<br>2.31 | 11.22;<br>1.98 | 12.21;<br>1.65 |
| Chr1 (C) | 11.22;<br>2.64 | 11.55;<br>1.65 | 11.55;<br>2.64 | 11.55;<br>1.65 | 11.55;<br>1.98 | 11.55;<br>1.65 | 11.55;<br>1.65 | 11.55;<br>1.32 | 11.55;<br>1.65 |
| Whole genome (W) | 87.13;<br>5.61 | 87.46;<br>3.96 | 88.45;<br>2.97 | 88.12;<br>3.96 | 88.45;<br>3.30 | - | 87.46;<br>3.96 | 88.45;<br>3.96 | 88.12;<br>3.96 |

#### Genomic features

Hypo found the maximum number of genomic features in each of the data-sets. For Illumina-only polished assemblies, the second best was amongst Racon or Pilon; Pilon did much better than Racon on Canu assemblies and Racon performed slightly better than Pilon on Wtdbg2 assembles. Unexpectedly, wtpoa-cns polishing, instead of increasing the number of features like other polishers, decreased them. The 2-round polishing using long reads slightly made the performance of wtpoa-cns2 better but it still lost the features that were present in the draft. Adding long reads to Hypo polishing slightly improved the results (an exception is Chr21 Canu assembly). Again, quite surprisingly, there was no change in Racon’s result with 2-round polishing.

#### BUSCO Score

There was no clear winner w.r.t BUSCO score. While wtpoa-cns produced the best score on some data-sets, it resulted in the worst score on others. On the whole genome, Racon produced the worst score. The score for Hypo-polished assembly, on each data-set was quite close to the best (if not the best).

## Tool availability

The tool and its source code are available on GitHub at the following link: https://github.com/kensung-lab/hypo

## Data source

Information about the sources of data-sets and other details have been listed in the Supplementary Material.

## Footnotes

i The commands used for various tools have been included in the supplementary material.

## References

Chin, C.-S., Alexander, D. H., Marks, P., Klammer, A. A., Drake, J., Heiner, C., Clum, A., Copeland, A., Huddleston, J., Eichler, E. E., Turner, S. W., and Korlach, J. (2013). Nonhybrid, finished microbial genome assemblies from long-read smrt sequencing data. Nature Methods, 10:563 EP –. Article.

Firtina, C., Bar-Joseph, Z., Alkan, C., and Cicek, A. (2018). Hercules: a profile HMM-based hybrid error correction algorithm for long reads. Nucleic Acids Research, 46(21):e125–e125.

Firtina, C., Kim, J. S., Alser, M., Senol Cali, D., Ercument Cicek, A., Alkan, C., and Mutlu, O. (2019). Apollo: A Sequencing-Technology-Independent, Scalable, and Accurate Assembly Polishing Algorithm. arXiv e-prints, page 1902.04341.

Fu, S., Wang, A., and Au, K. F. (2019). A comparative evaluation of hybrid error correction methods for error-prone long reads. Genome Biology, 20(1):26.

Jain, M., Koren, S., Miga, K. H., Quick, J., Rand, A. C., Sasani, T. A., Tyson, J. R., Beggs, A. D., Dilthey, A. T., Fiddes, I. T., Malla, S., Marriott, H., Nieto, T., O’Grady, J., Olsen, H. E., Pedersen, B. S., Rhie, A., Richardson, H., Quinlan, A. R., Snutch, T. P., Tee, L., Paten, B., Phillippy, A. M., Simpson, J. T., Loman, N. J., and Loose, M. (2018). Nanopore sequencing and assembly of a human genome with ultra-long reads. Nature Biotechnology, 36:338 EP –.

Koren, S., Walenz, B. P., Berlin, K., Miller, J. R., Bergman, N. H., and Phillippy, A. M. (2017). Canu: scalable and accurate long-read assembly via adaptive k-mer weighting and repeat separation. Genome Research, 27(5):722–736.

Laird Smith, M., Delany, N., Hepler, N., Alexander, D., Katzenstein, D., Brown, M., and Paxinos, E. (2016). An improved circular consensus algorithm with an application to detect hiv-1 drug resistance associated mutations (drams). In PacBio Conference Proceedings.

Lee, C. (2003). Generating consensus sequences from partial order multiple sequence alignment graphs. Bioinformatics, 19(8):999–1008.

Lee, C., Grasso, C., and Sharlow, M. F. (2002). Multiple sequence alignment using partial order graphs. Bioinformatics, 18(3):452–464.

Lee, H., Gurtowski, J., Yoo, S., Nattestad, M., Marcus, S., Goodwin, S., Richard McCombie, W., and Schatz, M. C. (2016). Third-generation sequencing and the future of genomics. bioRxiv.

Li, H. (2016). Minimap and miniasm: fast mapping and de novo assembly for noisy long sequences. Bioinformatics, 32(14):2103–2110.

Li, H. (2018). Minimap2: pairwise alignment for nucleotide sequences. Bioinformatics, 34(18):3094–3100.

Loman, N. J., Quick, J., and Simpson, J. T. (2015). A complete bacterial genome assembled de novo using only nanopore sequencing data. Nature Methods, 12:733 EP –.

Miga, K. H., Koren, S., Rhie, A., Vollger, M. R., Gershman, A., Bzikadze, A., Brooks, S., Howe, E., Porubsky, D., Logsdon, G. A., Schneider, V. A., Potapova, T., Wood, J., Chow, W., Armstrong, J., Fredrickson, J., Pak, E., Tigyi, K., Kremitzki, M., Markovic, C., Maduro, V., Dutra, A., Bouffard, G. G., Chang, A. M., Hansen, N. F., Thibaud-Nissen, F., Schmitt, A. D., Belton, J.-M., Selvaraj, S., Dennis, M. Y., Soto, D. C., Sahasrabudhe, R., Kaya, G., Quick, J., Loman, N. J., Holmes, N., Loose, M., Surti, U., Risques, R. a., Graves Lindsay, T. A., Fulton, R., Hall, I., Paten, B., Howe, K., Timp, W., Young, A., Mullikin, J. C., Pevzner, P. A., Gerton, J. L., Sullivan, B. A., Eichler, E. E., and Phillippy, A. M. (2019). Telomere-to-telomere assembly of a complete human x chromosome. bioRxiv.

Mikheenko, A., Prjibelski, A., Saveliev, V., Antipov, D., and Gurevich, A. (2018). Versatile genome assembly evaluation with QUAST-LG. Bioinformatics, 34(13):i142–i150.

Nanopore Technologies, O. (accessed June 2019). Medaka. https://nanoporetech.github.io/medaka/.

Roberts, M., Hayes, W., Hunt, B. R., Mount, S. M., and Yorke, J. A. (2004). Reducing storage requirements for biological sequence comparison. Bioinformatics, 20(18):3363–3369.

Roberts, R. J., Carneiro, M. O., and Schatz, M. C. (2013). The advantages of smrt sequencing. Genome Biology, 14(6):405.

Ruan, J. and Li, H. (2019). Fast and accurate long-read assembly with wtdbg2. bioRxiv.

Simão, F. A., Waterhouse, R. M., Ioannidis, P., Kriventseva, E. V., and Zdobnov, E. M. (2015). BUSCO: assessing genome assembly and annotation completeness with single-copy orthologs. Bioinformatics, 31(19):3210–3212.

Sović, I., Križanović, K., Skala, K., and Šikić, M. (2016). Evaluation of hybrid and non-hybrid methods for de novo assembly of nanopore reads. Bioinformatics, 32(17):2582–2589.

Vaser, R. and Šikić, M. (2019). Yet another de novo genome assembler. bioRxiv.

Vaser, R., Sović, I., Nagarajan, N., and Šikić, M. (2017). Fast and accurate de novo genome assembly from long uncorrected reads. Genome Research, 27(5):737–746.

Walker, B. J., Abeel, T., Shea, T., Priest, M., Abouelliel, A., Sakthikumar, S., Cuomo, C. A., Zeng, Q., Wortman, J., Young, S. K., and Earl, A. M. (2014). Pilon: An integrated tool for comprehensive microbial variant detection and genome assembly improvement. PLOS ONE, 9(11):1–14.

Warren, R. L., Coombe, L., Mohamadi, H., Zhang, J., Jaquish, B., Isabel, N., Jones, S. J. M., Bousquet, J., Bohlmann, J., and Birol, I. (2019). ntEdit: scalable genome sequence polishing. Bioinformatics.

Watson, M. and Warr, A. (2019). Errors in long-read assemblies can critically affect protein prediction. Nature Biotechnology, 37(2):124–126.

Weirather, J., de Cesare, M., Wang, Y., Piazza, P., Sebastiano, V., Wang, X., Buck, D., and Au, K. (2017). Comprehensive comparison of pacific biosciences and oxford nanopore technologies and their applications to transcriptome analysis [version 2; peer review: 2 approved]. F1000Research, 6(100).

Zhang, H., Jain, C., and Aluru, S. (2019). A comprehensive evaluation of long read error correction methods. bioRxiv.

Zook, J. M., Catoe, D., McDaniel, J., Vang, L., Spies, N., Sidow, A., Weng, Z., Liu, Y., Mason, C. E., Alexander, N., Henaff, E., McIntyre, A. B. R., Chandramohan, D., Chen, F., Jaeger, E., Moshrefi, A., Pham, K., Stedman, W., Liang, T., Saghbini, M., Dzakula, Z., Hastie, A., Cao, H., Deikus, G., Schadt, E., Sebra, R., Bashir, A., Truty, R. M., Chang, C. C., Gulbahce, N., Zhao, K., Ghosh, S., Hyland, F., Fu, Y., Chaisson, M., Xiao, C., Trow, J., Sherry, S. T., Zaranek, A. W., Ball, M., Bobe, J., Estep, P., Church, G. M., Marks, P., Kyriazopoulou-Panagiotopoulou, S., Zheng, G. X. Y., Schnall-Levin, M., Ordonez, H. S., Mudivarti, P. A., Giorda, K., Sheng, Y., Rypdal, K. B., and Salit, M. (2016). Extensive sequencing of seven human genomes to characterize benchmark reference materials. Scientific Data, 3(1):160025.

