## Supplementary Material for "HyPo: Super Fast & Accurate Polisher for Long Read Genome Assemblies"

### Hypo: Supplementary Material

#### Tools

##### POLISHERS

- Racon (v1.4.7)
- wtPoa-cns (Wtdbg2 0.0)
- ntEdit (v1.2)
- Pilon (pilon-1.23.jar) [only on chromosome-level data-sets]

##### OTHERS

- Minimap2 (v2.17-r941)
- Quast (v5.0.2)
- Samtools (v1.9)

#### Reference

- Reference: GRCh38.p13

#### Genomic Features

- GFF file

## HG002

- PacBio
  - Instrument:  PacBio RS II, Chemistry:  C3 & C4,Enzyme: P5 & P6
  - Coverage 69X
  - Read: BAM (and BAI):
- Illumina
  - 55X
  - Reads: BAM (and BAI)
- High Confidence Small Variants
  - Vcf (index):
  - Regions

### Commands

********** Direct Illumina Polishing **************

**MAPPING READS**

**Short reads**

minimap2 --MD -ax sr -t $NUMTH $DRAFT $R1 $R2 | samtools view -Sb - > mapped-sr.bam
samtools sort -@$NUMTH -o mapped-sr.sorted.bam mapped-sr.bam
samtools index mapped-sr.sorted.bam

**Long reads**

minimap2 --MD -ax $RTYPE -t $NUMTH $DRAFT $LONGR | samtools view -Sb - > mapped-lg.bam
samtools sort -@$NUMTH -o mapped-lg.sorted.bam mapped-lg.bam
samtools index mapped-lg.sorted.bam

[$RTYPE is map-pb or map-ont]

echo -e "$R1\n$R2" > il_names.txt
cd $CWD
#########################################################

**RACON**

minimap2 -t $NUMTH -x sr $DRAFT $READS  > $PREF.gfa.paf

racon -u -t $NUMTH -m 3 -x -5 -g -4 $READS $PREF.gfa.paf $DRAFT > $PREF.r.fa

#########################################################

**WTPOA-CNS**

samtools view $WORKDIR/mapped-sr.sorted.bam | wtpoa-cns -t $NUMTH -x sam-sr -d $DRAFT -i - -fo $PREF.w.fa

#########################################################

**HYPO**

$HYPO -d $DRAFT -r @$WORKDIR/il_names.txt -s $SZ -c $COV -b $WORKDIR/mapped-sr.sorted.bam \
-t $NUMTH -o $PREF.h.fa

########################################################
**NTEDIT**

PATH=$PATH:$NT/ntHits:$NT/ntEdit
$NT/ntEdit/ntedit-make ntedit draft=$DRAFT reads=$READS k=50 solid=true t=$NUMTH b=$PREF.n i=3 d=3 x=5 y=9

########################################################
**PILON**
java -Xmx128G -jar $PILON --genome $DRAFT --frags $WORKDIR/mapped-sr.sorted.bam \
--output $PREF.p --threads $NUMTH

********** Illumina + PacBio Polishing **************

########################################################
**RACON**

minimap2 -t $NUMTH -x $RTYPE $DRAFT $LONGR > $PREF.gfa1.paf

racon -u -t $NUMTH -m 3 -x -5 -g -4 $LONGR $PREF.gfa1.paf $DRAFT > $PREF.r1.fa

minimap2 -t $NUMTH -x sr $PREF.r1.fa $READS > $PREF.gfa2.paf

racon -u -t $NUMTH -m 3 -x -5 -g -4 $READS $PREF.gfa2.paf $PREF.r1.fa > $PREF.r2.fa
########################################################
**WTPOA-CNS**

samtools view $WORKDIR/mapped-lg.sorted.bam | wtpoa-cns -t $NUMTH -d $DRAFT -i - -fo $PREF.w1.fa

minimap2 --MD -ax sr -t $NUMTH $PREF.w1.fa $R1 $R2 | samtools view -Sb - > $PREF.sr.bam
samtools sort -@$NUMTH -o $PREF.sr.sorted.bam $PREF.sr.bam
samtools index $PREF.sr.sorted.bam

samtools view $PREF.sr.sorted.bam |wtpoa-cns -t $NUMTH -x sam-sr -d $PREF.w1.fa -i - -fo $PREF.w2.fa

########################################################

**HYPO**

$HYPO -d $DRAFT -r @$WORKDIR/il_names.txt -s $SZ -c $COV -b $WORKDIR/mapped-sr.sorted.bam -B $WORKDIR/mapped-lg.sorted.bam \
-t $NUMTH -p 96 -o $PREF.h2.fa

********** Quast **************

#####################################################

**CHR21 and CHR1**

quast -t 1 -o $STATSDIR -r $REF --eukaryote --conserved-genes-finding --min-identity 99.0 --no-sv --min-contig 1000 --no-read-stats --features $GFF -l $labels $files > $STATSDIR/out.txt

**Whole genome**

quast -t 1 -o $STATSDIR -r $REF --fragmented --large --conserved-genes-finding --min-identity 99.0 --no-sv --min-contig 1000 --no-read-stats --features $GFF -l $labels $files > $STATSDIR/out.txt
